# Layer 3 dynamically coordinates columnar activity according to spatial context

**DOI:** 10.1101/277533

**Authors:** Gijs Plomp, Ivan Larderet, Matilde Fiorini, Laura Busse

## Abstract

Spatial integration is a fundamental, context-dependent neural operation that involves extensive neural circuits across cortical layers of V1. To better understand how spatial integration is dynamically coordinated across layers we recorded single- and multi-unit activity and local field potentials across V1 layers of awake mice, and used dynamic Bayesian model comparisons to identify when laminar activity and inter-laminar functional interactions showed surround suppression, the hallmark of spatial integration. We found that surround suppression is strongest in layer 3 (L3) and L4 activity, showing rapidly sharpening receptive fields and increasing suppression strength. Importantly, we also found that specific directed functional connections were strongest for intermediate stimulus sizes and suppressed for larger ones, particularly for the L3->L5 and L3->L1 connections. Taken together, the results shed light on the different functional roles of cortical layers in spatial integration and show how L3 dynamically coordinates activity across a cortical column depending on spatial context.

## Introduction

One of the fundamental computations carried out by the primary visual cortex (V1) is the integration of visual information across space. In V1, neurons have a spatially localized classical receptive field (RF), but their activity strongly depends on the spatial context of the stimulus. Responses typically become larger for stimuli of increasing size, but can be suppressed if the stimulus extends beyond the RF (Knierim & van Essen 1992; Allman et al. 1985; Blakemore & Tobin 1972; DeAngelis et al. 1994; Gilbert & Wiesel 1990; Nelson & Frost 1978). This phenomenon, known as surround suppression, is thought to be a key mechanism for reducing redundancies in the natural input, perceptual pop-out and segmentation of object boundaries (Schmid & Victor 2014; Coen-Cagli et al. 2012; Sachdev et al. 2012; Angelucci et al. 2017).

Surround suppression is a hallmark of spatial integration that has been described at all stages of the retino-geniculo-cortical pathway, with dedicated mechanisms likely working in parallel (Angelucci et al. 2017). Starting at the level of retinal output, suppressive influences mediated by inhibition from amacrine cells with large RFs (Werblin 1972) can normalize responses of retinal ganglion cells (Solomon et al. 2006; Nolt et al. 2007; Alitto & Usrey 2008; Alitto & Usrey 2015). Compared to the retinal ganglion cell output, the strength of surround suppression increases in the lateral geniculate nucleus (dLGN) of the thalamus (Hubel & Wiesel 1961; Nolt et al. 2007). Because direct retinal input to dLGN is excitatory, this augmentation of surround suppression might be mediated by inhibitory influences within dLGN or from the thalamic reticular nucleus (Fisher et al. 2017), as well as by feedback provided from area V1 (Murphy & Sillito 1987; Nolt et al. 2007; Sillito & Jones 2002; Olsen et al. 2012). At the level of V1, further qualitative and quantitative changes in surround suppression have been observed: in supragranular layers of cats and primates, but much less so in thalamo-recipient layer 4 (L4), surround modulation is sharply orientation tuned (Hashemi-Nezhad & Lyon 2012; Henry et al. 2013a; Shushruth et al. 2013); in mice, cats and primates, RF size is smallest and surround suppression is strongest in supragranular layers (Self et al. 2014; Nienborg et al. 2013; Jones et al. 2000; Shushruth et al. 2009; Vaiceliunaite et al. 2013).

The local microcircuits in V1 that support surround suppression have been most extensively studied in L2/3. There, optogenetic studies in mice have revealed that one of the key circuits for surround suppression consists of L2/3 somatostatin-positive (SOM+) inhibitory interneurons, which are preferentially recruited by cortical horizontal axons (Adesnik et al. 2012). L2/3 SOM+ inhibitory interneurons have large RFs that effectively sum information across space while showing little surround suppression themselves. Furthermore, their inactivation results in decreased suppression of L2/3 principal cells (Adesnik et al. 2012). Consistent with a more general role in providing lateral inhibition, SOM+ neurons seem to control frequency tuning in L2/3 of mouse auditory cortex by providing lateral inhibition (Kato et al. 2017).

Besides horizontal connections and local inhibitory interneurons relevant for surround suppression in L2/3, additional mechanisms and circuits might control V1 spatial integration, potentially with differential impact across V1 layers. Surround suppression has been observed in most layers of V1, with varying RF sizes and suppression strengths (Nienborg et al. 2013; Self et al. 2014; Shushruth et al. 2009; Jones et al. 2000). In macaque L4C, for instance, extraclassical RFs are smallest, surround suppression is weakest and untuned for orientation, and emerges with response onset, consistent with L4C’s lack of long-range intracortical connections and driving input from the LGN (reviewed in Angelucci et al. 2017). In addition, V1 receives extensive interareal feedback connections, which preferentially terminate in L1 and L5/6 (Coogan & Burkhalter 1990; Markov et al. 2013) on both excitatory and local inhibitory interneurons (Gonchar & Burkhalter 2003; Zhang et al. 2014). These feedback connections can extend across multiples of the V1 RF diameter (Angelucci et al. 2002) and have been proposed to mediate suppressive influences from surround regions further away (Angelucci et al. 2002; Angelucci & Bressloff 2006). While each of these mechanisms and neural circuits likely contributes to surround suppression across several V1 layers, the coordination of suppression across V1 layers remains poorly understood.

To shed light on how activity across layers of a V1 column is differentially orchestrated during spatial integration, we characterized with high temporal resolution surround-suppressed activity within each layer, and surround-suppressed functional connectivity between layers. Directed functional connectivity analysis in the Granger-causality framework (Granger 1969; Seth et al. 2015) has previously provided informative models of how cortical layers or areas interact in sensory processing (Michalareas et al. 2016; Saalmann et al. 2012; Liang et al. 2017; van Kerkoerle et al. 2014). In awake mice, we recorded single- and multi-unit activity as well as local field potentials (LFPs) across all six layers, computed dynamic functional connectivity estimates (Milde et al. 2010; Baccalá & Sameshima 2001; Plomp, Quairiaux, Michel, et al. 2014) and used a novel Bayesian model comparison approach to identify at what latencies surround suppression was evident in laminar activity and in inter-laminar functional connectivity strengths. We found sustained surround suppression at L4 and L3, with a rapid sharpening of the tuning profile at early latencies after stimulus onset. L3 also showed persistent surround-suppressed inter-laminar connectivity that specifically influenced L1 and L5 at early latencies. L4, however, did not show such persistent surround-suppressed connections. Together, these results demonstrate a key role of L3 in orchestrating activity across layers of V1 in a size-dependent way.

## Methods

### Experimental procedures

All experiments were performed on awake, adult mice. The procedures complied with the European Communities Council Directive 2010/63/EC, the German Law for Protection of Animals, and were approved by local authorities following appropriate ethics review.

### Mice

We used 7 adult mice (2 C57BL/6J and 5 mice with floxed NR1 receptors used as controls for a different study (see Korotkova et al. 2010 for details on the mouse line); 4 males, 3 females), which ranged in age from 2 - 7 months. We used recordings with at least two contacts both in L1 and L6 (26 experiments, from 13 penetrations), allowing bipolar derivation for L1 to L6 (see below).

### Surgical procedures

Surgeries were performed as described previously (Erisken et al. 2014). Briefly, mice were anesthetized using 3% Isoflurane, which was maintained for the duration of the surgery at 1.5-2%. Analgesics (Buprenorphine, 0.1 mg/kg, sc) was administered, and eyes were prevented from dehydration with an ointment (Bepanthen). The animal’s temperature was kept at 37°C via a feedback-controlled heating pad (WPI). A custom-designed head post was attached to the anterior part of the skull using dental cement (Tetric EvoFlow, Ivoclar Vivadent), and two miniature screws were placed in the bone over the cerebellum, serving as reference and ground (#00-96X 158 1/16, Bilaney). Following the surgery, antibiotics (Baytril, 5mg/kg, sc) and long-lasting analgesics (Carprofen, 5mg/kg, sc) were administered for 3 consecutive days. After recovery, mice were placed on a Styrofoam ball and habituated to head-fixation for several days. The day before electrophysiological recordings, mice were again anesthetized (Isoflurane 2%), and a craniotomy (~1 mm^2^) was performed over V1 (3 mm lateral from the midline suture, 1.1 mm anterior to the transverse sinus). The exposed brain was sealed with the silicon elastomer Kwik-Cast at the end of each recording session. Recording sessions always started at least one day after surgery.

### Visual stimuli

Visual stimuli were created with custom software (Expo, https://sites.google.com/a/nyu.edu/expo/home), and presented on a gamma-corrected LCD monitor (Samsung 2233RZ; mean luminance 50 cd/m^2^) placed 25 cm from the animal’s eyes.

To measure RFs, we mapped the ON and OFF subfields with a sparse noise stimulus. The stimulus consisted of white and black squares (4° diameter) briefly flashed for 150 ms on a square grid (40° diameter). To measure size tuning we centered circular square-wave gratings (spatial frequency 0.02 cyc/deg) of 10 different diameters (3.9, 5.6, 7.8, 12.1, 15.5, 21.8, 30.6, 43.1, 60.5 or 67.3° of visual angle) on online estimates of RF centers based on threshold crossings, and presented each stimulus in pseudo-random order for 750 ms, followed by a 500 ms ISI. We also included a blank screen condition, in which only the mean luminance gray screen was presented. Orientation of the gratings was chosen to match the average preferred orientation tuning based on threshold crossings. The number of trials for each stimulus size varied across animals (mean 208, range 50 - 500). For determining the L4/L5 border using current source density analysis, we presented a full-field, contrast-reversing checkerboard at 100% contrast, with a spatial frequency of 0.02 cyc/deg and a temporal frequency of 0.5 cyc/s.

### Extracellular recordings

Extracellular recordings were performed in awake head-fixed mice placed on a Styrofoam ball. Recordings of neural activity were performed with a 32 channel linear silicon probe with 25 μm inter-contact spacing (Neuronexus, A1×32-5mm-25-177-A32).

Extracellular signals were recorded at 30 kHz (Blackrock microsystems) and analyzed with the NDManager software suite (Hazan et al. 2006). For spike sorting, we divided the linear array into 5 “octrodes” (8 channels per group with 2 channels overlap). Using a robust spike detection threshold (Quiroga et al. 2004) set to 6 SDs of the background noise, we extracted spike-waveshapes from the high-pass filtered continuous signal. The first 3 principal components of each channel were used for automatic clustering with a Gaussian Mixture Model in KlustaKwik (Henze et al. 2000), and the resulting clusters were manually refined with Klusters (Hazan et al. 2006). Duplicate spike clusters, which can arise from separating the electrode channels in different groups for sorting, were defined as pairs of neurons, for which the cross-correlogram’s zero-bin was 3 times larger than the mean of non-zero bins, and one of the neurons in the pair was removed from the analysis.

For calculating the envelope of multi-unit activity (MUAe), we full-wave rectified the median-subtracted, high-pass filtered signals, before low-pass filtering (200 Hz) and down-sampling to 2000 Hz (Self et al. 2014; Supèr & Roelfsema 2005; van der Togt et al. 2005). To assure spatial alignment of RFs across cortical depth, we routinely assessed RF maps obtained by the sparse noise stimulus, for which we used average MUAe between 50 and 175 ms after stimulus onset (for an example, see Fig. 1b). For analysis of size tuning, MUAe was normalized to pre-stimulus values for each single trial and layer. Response onset (Figure 2) was quantified as the first latency after which the 95% CI across animals remained above 0 for at least 10 ms, for the largest presented grating.

**Figure 1.**
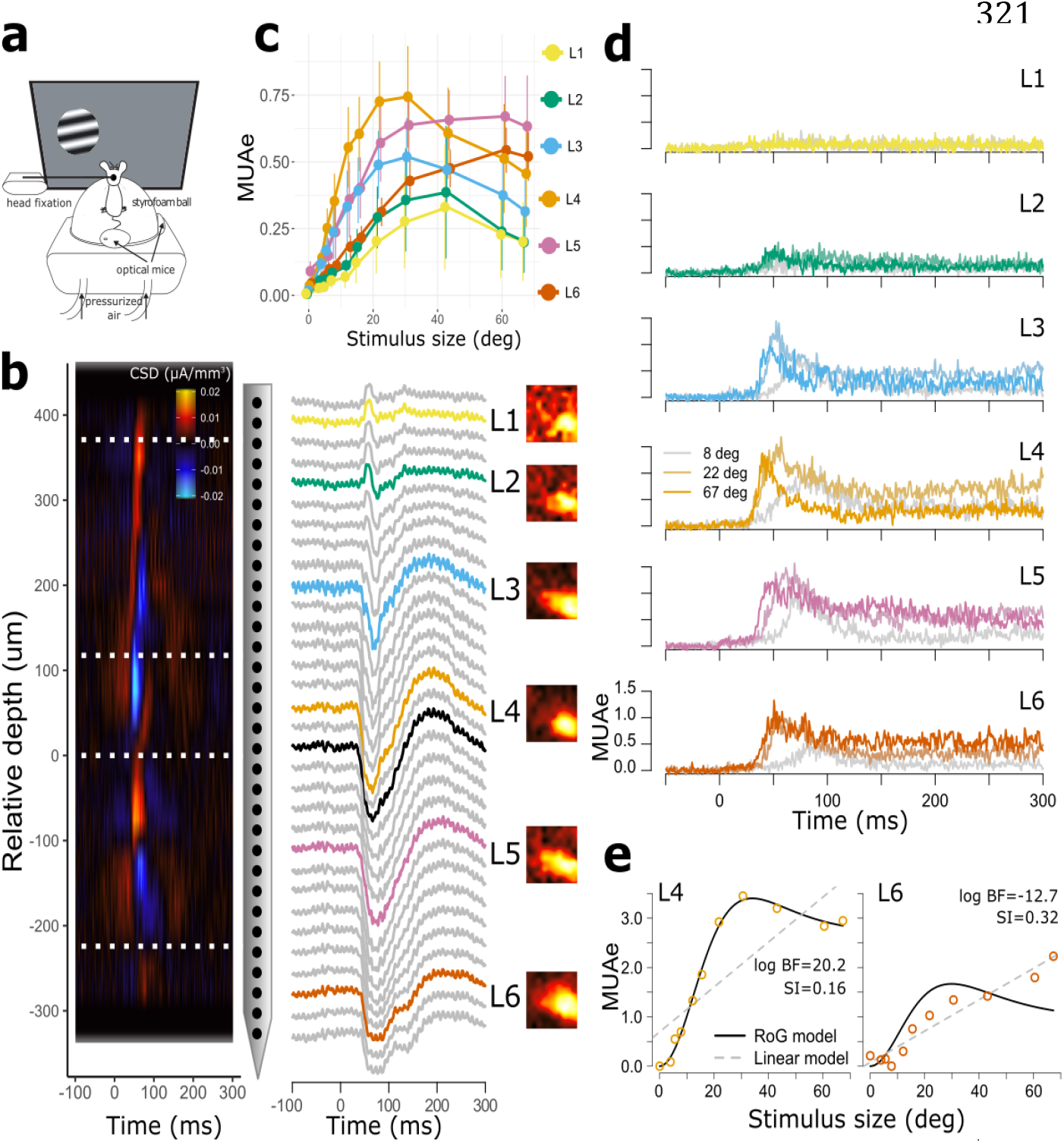
Size tuning across V1 layers. **a**) Recording setup. **b**) Left: CSD pattern evoked by a contrast-reversing checkerboard stimulus for an example penetration across V1 cortical depth reveals base of L4. Dashed lines indicate layer boundaries based on histology (Heumann et al. 1977). Middle: corresponding LFP signal with labeling of putative cortical layers. Black: LFP responses at the contact closest to the base of L4; color: LFP responses from the middle of each layer, which were used for further analyses. Right: MUAe receptive field (RF) maps for the same penetration measured using a sparse noise stimulus, showing consistent RF locations across electrode depth. **c**) Median MUAe across time as a function of stimulus diameter. Error bars denote standard errors across animals (n = 7 mice). **d**) Evoked grand-average MUAe for three grating diameters in L1 through L6. **e**) Illustration of competing linear and RoG model fits to individual MUAe of L4 and L6 at 48 ms after stimulus onset, with positive evidence in favor of the RoG model for L4 but not L6.

**Figure 2.**
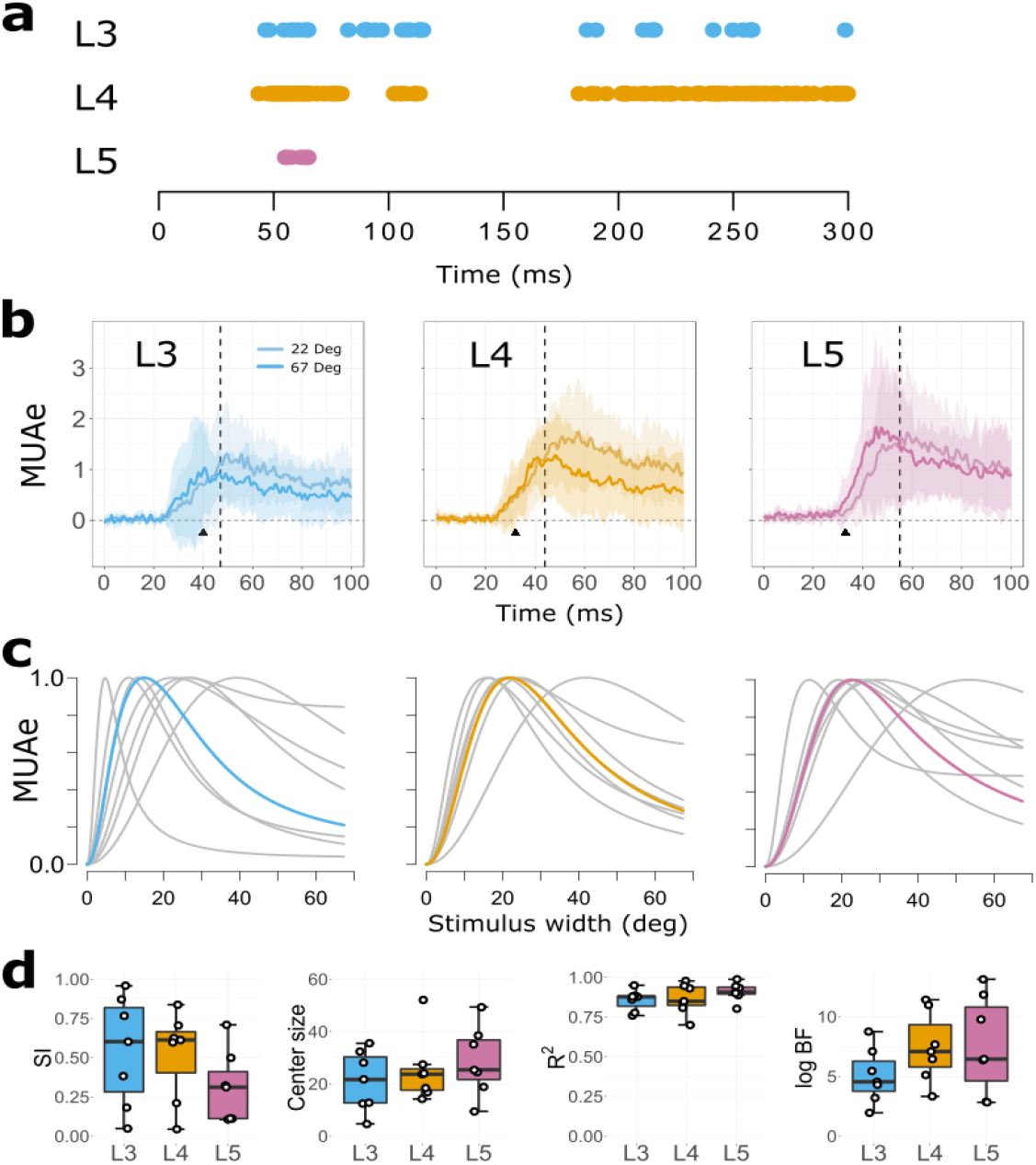
Dynamics of multi-unit size tuning in L3, L4 and L5. **a**) Time points after stimulus onset where MUAe showed a surround-suppressed tuning curve profile in at least 6/7 animals. **b**) Mean MUAe across animals (n=7) relative to stimulus onset for 22° (light colors) and 67° gratings (dark colors), for L3 (right), L4 (middle) and L5 (right). The black vertical lines indicate indicate onsets of surround suppression, response onset is indicated with a black triangle (see Methods). Shading reflects 95% CI across animals. **c**) Median tuning curves for stimulus size (colored), averaged across time points with significant suppression (see a); grey curves are fits from individual animals. **d**) Median surround suppression indices (SI), center sizes, R^2^ and log B_12_ values across time points.

The LFP was computed by downsampling the data to 1250 Hz, and high pass, forward-backward filtering at 1Hz (2^nd^ order Butterworth). To map electrode contacts to cortical layers, we computed current source density (CSD) from the second spatial derivative of the LFP (Mitzdorf 1985) and assigned the base of L4 to the contact that was closest to the earliest CSD polarity inversion from sink to source. The remaining contacts were assigned putative layer labels based on the known relative thickness of V1 layers (Heumann et al. 1977), and an assumed total thickness of ~1 mm. We checked the L4-L5 boundary localization using CSD methods that do not assume constant activity in the horizontal direction (Pettersen et al. 2006), and obtained identical depth estimates.

We then selected for further analysis the channel closest to the middle of each layer as the representative signal for that layer. To reduce volume conduction effects from neural and noise sources (e.g., muscles artifacts) we derived bipolar LFPs by subtracting signals from the two neighboring electrodes (Trongnetrpunya et al. 2015; Rohenkohl et al. 2018; Bastos et al. 2015). Power spectral density (PSD) was calculated with the S-transform on epochs of −500 to 500 ms around stimulus onset and rectified as relative increases with respect to prestimulus activity (Roberts et al. 2013).

### Time-varying directed connectivity

Functional connectivity values were calculated from single trial bipolar LFP signals between −50 to 300 ms after stimulus onset. We chose bipolar LFPs (Bastos et al. 2015; Rohenkohl et al. 2018; Trongnetrpunya et al. 2015) because possible noise amplification in CSD calculations can negatively impact connectivity analysis (Trongnetrpunya et al. 2015). We used a time-varying implementation of the Partial Directed Coherence (PDC, Baccalá & Sameshima 2001), a multivariate, directed connectivity measure based on the notion of Granger causality, or the relative predictability of signals from one another (Granger 1969; Bressler & Seth 2011).

PDC was derived from a multivariate autoregressive model of the recorded signals, which is based on a fixed model order that reflects the maximum time lag of observations included in the model (Baccalá & Sameshima 2001). Optimal model orders were determined by minimizing Akaike’s information criterion across epochs within animals for each stimulus size (Barnett & Seth 2014), and ranged between 13 and 15 (10-12 ms). This parametric approach avoids known pitfalls of some non-parametric approaches (Stokes & Purdon 2017).

To obtain time-varying multivariate autoregressive (tvMVAR) models we used a Kalman filter approach (Milde et al. 2010). The constants that determine adaptation speed during parameter estimation were fixed at 0.02, following previous work (Plomp, Quairiaux, Michel, et al. 2014; Plomp, Quairiaux, Kiss, et al. 2014; Astolfi et al. 2008). Within animals, tvMVAR parameter estimates were averaged across trials for the 11 conditions (Ghumare et al. 2015).

We orthogonalized the tvMVAR parameters to further guard against possible volume conduction effects (Omidvarnia et al. 2014; Hipp et al. 2011), and obtained PDC values using a row-wise normalization to optimize sensitivity to information outflows (Astolfi et al. 2007; Kuś et al. 2004):

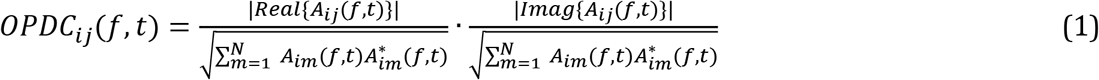

where A is the frequency-transformed tvMVAR parameter matrix. We squared OPDC values to enhance accuracy and stability (Astolfi et al. 2006). Resulting PDC matrices were normalized (0-1) for each animal across conditions, time and frequencies (1-150 Hz) and multiplied by the normalized spectral power across conditions, time and frequencies, obtaining a weighted PDC (wPDC) estimator that has been shown to better reflect the underlying physiological processes (Plomp, Quairiaux, Michel, et al. 2014).

### Bayesian model comparison

In V1, the suppressive influence from the extraclassical surround is generally considered a phenomenon accounted for by divisive normalization (Carandini & Heeger 2011). On a descriptive level, effects of surround suppression in spatial tuning can be captured by a Ratio of Gaussians (RoG) model (Cavanaugh et al. 2002a), where a center Gaussian with independent amplitude and width is normalized by a Gaussian representing the surround. Thus, responses are given by:

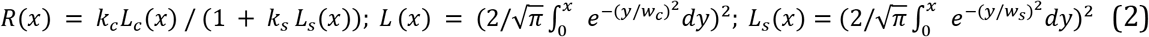

where *x* is the stimulus diameter, *k_c_* and *k_s_* are the gains of center and surround, *w_c_* and *w_s_* their respective spatial extents, and *L_c_* and *L_s_* are the summed squared activities of the center and surround mechanisms, respectively. Our use of the ROG model is not meant to reflect a particular biophysical implementation, but should only serve as a quantitative description of tuning; yet, it has been shown that the RoG model applied to V1 responses can outperform models assuming subtractive influences from the surround (Cavanaugh et al. 2002b). While RoG models can also capture non-suppressed responses, responses increasing monotonically for most of the tested stimulus sizes can be more parsimoniously explained by a simple linear null model with only two parameters:

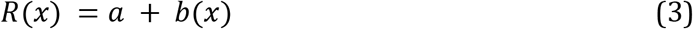

where *a* and *b* reflect intercept and slope respectively. In Bayesian statistics the evidence in favor of one model (M_1_; RoG model) over another (M_2_; linear null model) given the data is the ratio of their posterior probabilities, or Bayes factor (Raftery 1995; Rouder et al. 2009; Jeffreys 1998):

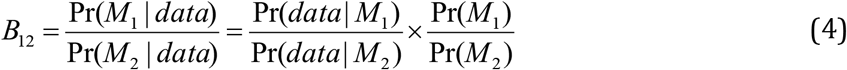

We assumed both models to be equally likely a priori, and set the summed prior probabilities to 1. Bayes factors were approximated using the Bayesian Information Criterion (BIC) values associated with M_1_ and M_2_ (Raftery 1995):

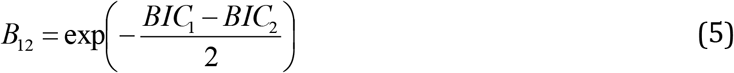

Model comparisons based on BIC values penalize for the number of parameters and here provide a conservative approach for detecting evidence in favour of the RoG model. The Bayes factor (B_12_) quantifies the relative amount of evidence in the data for each model. B_12_ > 3 is generally considered positive evidence in favor of M_1_ (Kass & Raftery 1995; Raftery 1995). We report B12 values on logarithmic scales.

For each timepoint in the MUAe activity, and for each time-frequency point in the connectivity analysis, we fitted amplitudes as a function of stimulus size with linear and RoG models (non-linear least squares, Port algorithm), enforcing *w_c_* < *w_s_ (Cavanaugh et al. 2002b).* Model comparisons were done separately for each animal to avoid effects driven by single animals or outliers.

From the RoG models we obtained RF center size as the stimulus diameter eliciting peak amplitude (MUAe or connectivity strength); models with center sizes below 3.9° (smallest presented size) were not further analyzed. We quantified strength of suppression using the suppression index *(SI):*

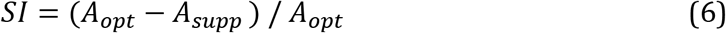

where A is MUAe, spike rate or wPDC amplitude, *A_opt_* is the model’s peak amplitude and *A_supp_* is the amplitude at the largest presented size (67.3°) (DeAngelis et al. 1994; Self et al. 2014).

We identified data points with a suppressed tuning curve profile as those that showed both positive evidence in favor of the RoG model (B_12_ > 3, or equivalently log B_12_ > 1), and a nonzero suppression index (SI > 0). This latter requirement ensured that responses with asymptotic or other non-linear monotonic increases were not further considered. For further analysis, we retained data points where at least 6/7 animals passed both criteria (conjunction analysis). When an animal failed a criterion, its results were not included for further summaries. Layers or functional connections with only one data point were not further analyzed (L6 MUAe at 288 ms; L2->L1 connection at 82 ms, 6Hz). All model comparisons and analyses were done in R (www.r-project.org). For graph visualization (Figure 6), the layout was determined using the Fruchterman-Reingold algorithm (Fruchterman & Reingold 1991), applied to a binary adjacency matrix, as implemented in the igraph library for R.

For the model comparison analysis of single-unit RF dynamics, we included sorted units from the central contact in the target layer and the two contacts immediately above and below (i.e. across 5 contacts, covering 125 μm centered around the middle of L3 or L4, Figure 1b). We first fit RoG models to the spike rates between 0 and 300 ms after stimulus onset to identify units with R^2^ > 0.5, SI > 0 and center size > 4°. For these units we then dynamically fit RoG models in 50 ms bins sliding between 25 and 250 ms after stimulus onset (1 ms shift size).

## Results

In awake, head-fixed mice (Figure 1a), we performed extracellular recordings across all layers of area V1 (Figure 1b). We assigned electrode contacts to layers based on CSD analysis (Mitzdorf 1985) (Figure 1b), and computed from the recorded signals the local field potentials (LFP) and the envelope of multi-unit activity (MUAe) (Supèr & Roelfsema 2005; van der Togt et al. 2005). MUAe responses reflect the number and amplitude of spikes close to the electrode, resembling thresholded multi-unit data and average single-unit activity (Self et al. 2014; Supèr & Roelfsema 2005). To assess spatial integration, we presented gratings of various diameters centered on the RFs of the recorded neurons. Similar to numerous studies before, we found that time-averaged multi-unit activity at the granular and supragranular layers varied systematically with grating diameter, typically peaking at intermediate sizes of around 20-30° of visual angle and showing surround suppression with larger diameters (Figure 1c). Beyond these time-averaged response patterns, we observed considerable variation in response latencies, amplitudes and time course of MUAe responses across layers depending on stimulus size (Figure 1d).

To get first insights into the dynamics of size tuning at each layer, we used Bayesian model comparisons to identify at what latencies MUAe activity showed more evidence in favor of a RoG model than a linear model (Figure 1e). The RoG models consist of two Gaussians with the same center location but different widths and amplitudes, and can well capture V1 size tuning curves (Cavanaugh et al. 2002b; Vaiceliunaite et al. 2013). In this model, preferred center size is given by the peak location of the fitted curve and suppression strength is the amplitude reduction for large stimuli relative to peak response (suppression index SI, Van den Bergh et al. 2010). By selecting RoG models with B12 values > 3 and SI > 0, we identified layers and time points where MUAe amplitudes consistently reflected a surround-suppressed tuning curve (see Methods, Bayesian model comparison).

### Multi-unit activity in layers 3-5 is dynamically suppressed

Using the above outlined, stringent model comparison approach, we found that the major time points where MUAe activity consistently reflected surround suppressed tuning curves occurred in L3, L4, and L5 (Figure 2a). Surround suppression was first evident in L4, emerging at 44 ms after stimulus onset (first data point with consistent evidence in favor of suppression in 6/7 animals). L4 suppression onset came 12 ms after response onset at 32 ms (first data point after which the 95% CI across animals exceeded baseline for at least 10 ms; Figure 2b, middle). The observed short delay between stimulus-driven activity and surround suppression onset is consistent with the hypothesis that even at the earliest latencies in L4 of mouse V1, surround suppression is not solely inherited from the dLGN, but is rapidly shaped by intracortical circuits (Knierim & van Essen 1992; Smith et al. 2006). L4 MUAe suppression continued until 114 ms after stimulus onset, and also showed a later, sustained surround-suppressed response component, from 183 ms onward (Figure 2a). Across timepoints, L4 MUAe peaked for stimulus diameters of 24° (range: 14 – 52°), with a median SI of 0.61 (range: 0.04 – 0.84) (Figure 2c, d).

In L3, MUAe onset (40 ms) occurred later than in L4, but the onset of suppression was similar to that of L4 (47 ms; Figure 2b), and overall, surround suppression had a similar time course and strength (Figure 2a, c, d). In particular, L3 MUAe showed surround suppression during broadly two periods: an early period starting slightly after response onset from 47 to 115 ms, and a later one between 187 and 300 ms (end of epoch; Figure 2a). Overall, L3 MUAe median center size was 22° (range: 5 - 36°) with suppression strengths (SI) of 0.6 (0.05 - 0.96) (Figure 2c, left), indicating considerable suppression of RF activity in L3 during surround stimulation.

In L5, surround suppression started relatively late, at 55 ms after stimulus onset, or 22 ms after response onset (at 33 ms, Figure 2a, b), and suppression was more concentrated in time (55 – 65 ms) than in the more superficial layers. At these relatively few time points, however, consistent evidence for surround suppression was found in all mice. Here, L5 MUAe preferred stimulus diameters of 25° (range: 9 – 49°), and a median SI of 0.31 (0.11–0.71). Model comparison results and RoG model parameters for the surround suppressed MUAe per layer are summarized in Table 1.

**Table 1.**
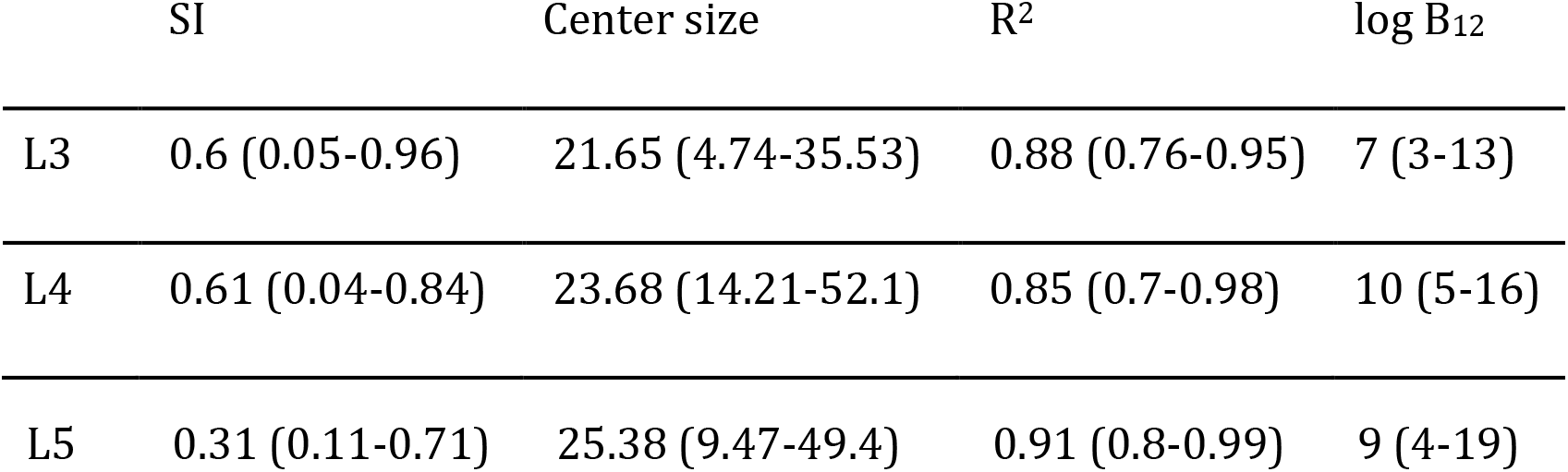
RoG model profiles for surround-suppressed MUAe activity per layer, median (min, max) across mice.

None of the MUAe in L1 and L6 showed strong and consistent evidence for surround suppression at any single time-point. In L1, stimulus-evoked transient MUAe responses were small and showed only modest variations with stimulus size across time (Figure 1d). This is not surprising, given that L1 has relatively few neurons (Hestrin & Armstrong 1996; Gonchar et al. 2007), whose spiking activity is difficult to pick up with extracellular recordings. In L6, by contrast, stimulus-evoked MUAe was strong, but it increased monotonically with stimulus size without showing consistent suppression at any single time point (Figure 1c,d). The weak average SI in L5 and the relative lack of surround suppression in L6 are in line with previous studies, which reported broader spatial tuning for V1 infragranular layers in cats (Jones et al. 2000), and mice (Self et al. 2014; Nienborg et al. 2013; Vaiceliunaite et al. 2013).

### Functional connectivity between layers shows size tuning

Having observed strikingly different effects of surround suppression across cortical layers and in time, we next assessed how cortical layers dynamically orchestrate activity during spatial integration by analyzing inter-laminar functional connectivity. We calculated time-varying connectivity between all layers based on Partial Directed Coherence (PDC), a multivariate variant of Granger causality in the frequency domain (Baccalá & Sameshima 2001; Bressler & Seth 2011; Plomp, Quairiaux, Michel, et al. 2014; Seth et al. 2015) (see also Methods, Time-varying directed connectivity). Granger causality is a statistical measure of time-lagged regularities between recorded signals (here LFPs with bipolar derivation), where increased connectivity means that the future activity of the target layer becomes better predictable from the activity at the source layer, i.e., that the source layer more strongly drives activity in the target layer. Since driving in this functional connectivity framework might or might not occur via direct anatomical connectivity, we decided to interpret our results in light of the simultaneously recorded spiking activity (MUAe) and the known structural connectivity, while also emphasizing alternative explanations and limits of the technique (see also Discussion). In the past, similar connectivity analyses have helped to better understand laminar interactions at rest and during sensory processing (Bollimunta et al. 2008; van Kerkoerle et al. 2014; Plomp, Quairiaux, Kiss, et al. 2014; Chen et al. 2017; Liang et al. 2017).

We first applied the analysis of PDC to our V1 laminar recordings, and asked how directed functional connectivity depended on spatial context. We reasoned that, in the same way that surround-suppressed activity reflects contextual influences on neuronal responsivity, surround-suppressed connections would reflect how stimulus context parametrically varies the influence that a source layer has on future activity of its target layer. We illustrate this reasoning in Figure 3, using two example connections. Figure 3a shows directed connectivity strengths in response to a large-sized grating from L4 to each of the other layers (for other source layers, see Supplementary Figure 1). In line with known L4 projections, the main targets of L4 driving were L3 and L5 (Xu et al. 2016; Pluta et al. 2015; Thomson 2003; Harris & Shepherd 2015). Figure 3b shows connectivity strengths of the L3 -> L1 connection, for different stimulus sizes, and highlights that connection strength can depend on spatial context.

**Figure 3.**
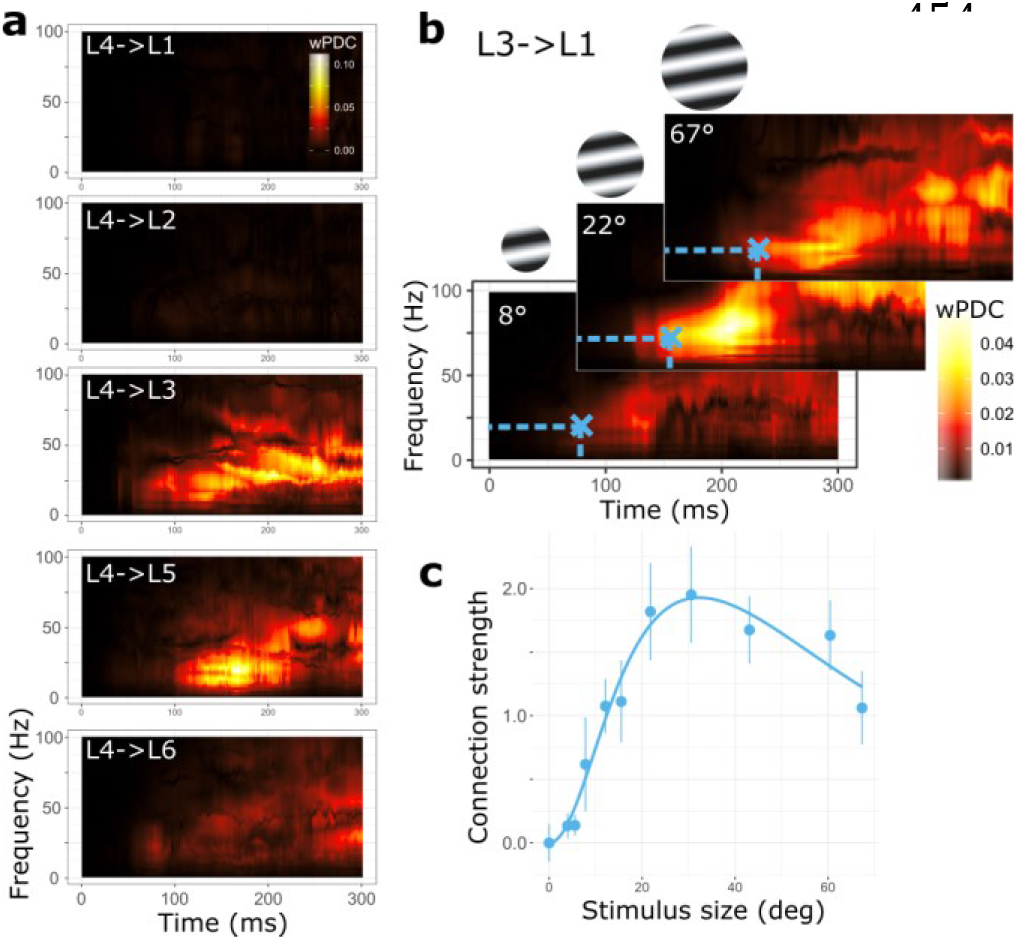
Inter-laminar functional connectivity. **a**) Time-frequency plots of directed functional connectivity strengths (median across animals) from L4 to the other layers for a 67° grating. **b**) Median L3 to L1 connectivity strengths for three grating sizes, exhibiting a surround-suppressed tuning-curve at the indicated time-frequency point (blue cross). **c**) RoG model of the average connectivity strengths across animals as a function of stimulus size, for the time-frequency point indicated by the blue cross in **b**). Error bars denote s.e. around the mean, n=7.

Applying the same Bayesian model comparison approach as above to directed functional connectivity strengths at each time (0-300 ms) and frequency point (1-150 Hz), we obtained for each connection a time-frequency distribution of Bayes factors (B12). Using identical thresholding and conjunction analysis (see Supplementary Figure 2 for all size-tuned connections before conjunction analysis) as for MUAe, we identified five major inter-laminar connections whose strength followed a surround-suppressed tuning curve and hence relayed information about stimulus size and context.

The earliest surround-suppressed connection extended from L4 to L2 at latencies between 49 and 51 ms after stimulus onset, and operated in the beta band (Figure 4a). Across time and frequency points the L4->L2 connection was strongest for stimuli of 23° (range across animals: 10-41°) and strongly suppressed for larger stimuli (median SI = 0.65, range 0.05-0.89). In general, functional connectivity from L4 to L2 is consistent with the known ascending projections from L4 to L2/3 (Xu et al. 2016; Thomson 2003; Harris & Shepherd 2015; Pluta et al. 2017). The surround-suppressed L4->L2 connectivity coincided with the onset of surround-suppressed MUAe at L4 (Figure 2a), consistent with the notion that size-tuned multi-unit activity plays a role in the relay of size information to L2. Remarkably,-the time point by time point MUAe analysis of L2, did not show consistent evidence for surround-suppressed activity. This indicates that the L4->L2 driving does not immediately and consistently result in size-tuned MUAe at L2. Instead, it suggests that L4 activity parametrically drives postsynaptic potentials at L2 in a less time-locked manner, contributing to the surround-suppressed activity obtained in time-averaged data (Figure 1c).

**Figure 4.**
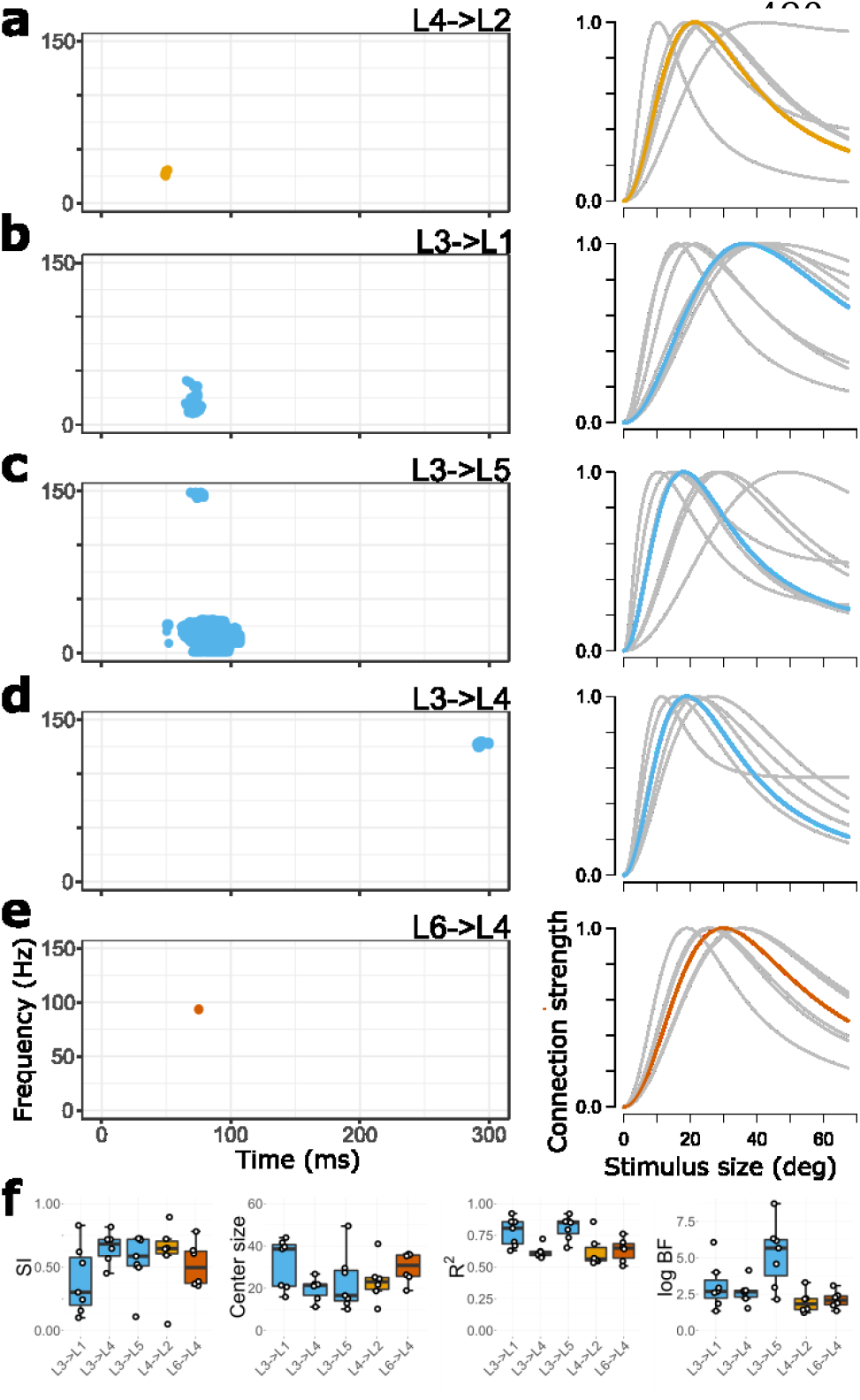
Functional connections relaying information about stimulus size and surround context. **a-e**) Time-frequency points where functional connections (wPDC) for at least 6/7 animals showed positive evidence for surround-suppression. Plots on the right show the corresponding median size tuning curve (colored) and individual animal data (grey), averaged across the colored regions in (a-e). f) Box and whisker plot of suppression strength (SI), preferred size, R^2^ and log B_12_ values for each surround-suppressed connection. Color coding is for source layer, corresponding to Figure 1.

A second ascending connection with a surround-suppressed driving profile was the L3->L1 connection, which showed surround-suppressed connectivity in the beta and low gamma band between 65 and 78 ms (Figure 4b). For this connection median preferred size was 39° (16-44°) with a suppression strength of SI = 0.3 (0.1 - 0.83) (Figure 4d). As with the L4->L2 connection, the latencies of surround-suppressed driving coincided with a period of surround-suppressed MUAe in the source layer, and with an absence of evidence for time-resolved surround suppression in the target layer (Figure 2d), suggesting that size-tuned activity at L3 relays size information to L1 by driving postsynaptic potentials with low temporal precision. Target L1 is an important recipient of thalamic and cortical feedback (D’Souza & Burkhalter 2017; Ji et al. 2015; Coogan & Burkhalter 1993). L1 postsynaptic potentials, in turn, have modulatory influence throughout the column because neurons in most layers have apical dendrites in L1, allowing L1 to change spike likelihoods in deeper layers (Egger et al. 2015; Jiang et al. 2013; Larkum et al. 1999). The L3->L1 driving was largest for stimuli inside the RF of the column, indicating that the processing at L1 is modulated in a size and context-dependent way. Hence, the L3->L1 connection can potentially modulate how feedback arriving at L1 affects activity throughout the column.

Besides these ascending size-tuned connections, size information was also relayed via descending functional connectivity from L3. Driving from L3->L5 showed a surround-suppressed profile across several frequency bands at latencies between 50 and 107 ms (Figure 4c). Functional connectivity was overall strongest for stimuli spanning 17° (range: 10-50°), with a suppression strength of 0.59 (SI, range 0.11-0.73). L2/3 pyramidal cells constitute the main input to L5, and L3 contains apical dendrites from L5 pyramidal cells (Thomson & Bannister 2003; Xu et al. 2016). Functional synaptic coupling between L3 and L5 has been previously established using laminar population analysis (Einevoll et al. 2007). The size-tuned L3->L5 connection coincided with size-tuned MUAe at L3, in line with the idea that spiking activity in L3 drives postsynaptic potentials at L5 in a context-dependent manner. At the target layer L5, size-tuned MUAe coincided with this connection, indicating that the surround-suppressed L3->L5 connection may immediately contribute to surround-suppressed spiking activity at L5. L5 too is an important recipient of feedback connections (Coogan & Burkhalter 1990; Markov et al. 2013), suggesting the possibility that this L3 driving modulates the influence of input from other areas on processing in this column.

At longer latencies, between 290 and 300 ms (end of epoch), the L3->L4 connection showed surround suppression in the high gamma band (Figure 4d), in line with known excitatory projections (Xu et al. 2016). The median center size of this connection was 21° (range: 11 - 27°) with an SI of 0.68 (0.45-0.82). This timing corresponds to the second period of size-tuned MUAe in L4. While our results are thus consistent with the hypothesis that late L3->L4 driving shapes L4 size-tuned activity, it is likely that other influences at these latencies also contribute to L4 surround suppression.

The relays of surround-suppressed size information from L3 to both L5 and L1, and later to L4, thus all occurred simultaneously with L3 surround-suppressed MUAe (Figure 2). The coexistence of surround-suppressed spiking activity and surround-suppressed driving from L3 is consistent with the interpretation that surround-suppressed population activity at L3 has a major role in orchestrating activity across V1 layers in a context-dependent way.

Lastly, the ascending L6->L4 connection briefly showed surround-suppressed size tuning in the gamma band, with peak driving for stimuli of 31° (range: 19 - 36°) and median SI of 0.5 (0.35-0.78; Figure 4e). Occurring at 75 ms, considerably later than V1 response onset, this functional connection might be driven by fast feedback from higher-level areas to L6 (Domenici et al. 1995; Nowak et al. 1997; Zhang et al. 2014). Simultaneously, size-tuned MUAe was seen at target layer L4, suggesting that L6 driving contributes to size-tuned activity at L4 at those latencies. The surround-suppressed L6->L4 connectivity might enhance the gain of visual input in L4 through intracortical circuits (Raizada & Grossberg 2003). In our data, however, the mechanism of this driving remains unclear because L6 MUAe did not show surround suppression.

Taken together, we found that directed functional connectivity strengths from L3, L4 and L6 resemble surround-suppressed tuning curves that are typically observed for single-units. These connections most strongly influenced target layers for stimuli covering the RF and showed reduced driving for larger stimuli. L3 in particular, but also L4 and L6, thus effectively relay information about stimulus size, and play an active role in coordinating laminar activity patterns through parametric variations in connection strengths that depend on spatial context.

### Dynamic size tuning

Previous work has shown that spatial RF properties in V1 can undergo fast dynamics after response onset, showing rapid decreases in preferred size and increases in suppression in both cat and monkey (Briggs & Usrey 2011; Malone et al. 2007; Wörgötter et al. 1998). We therefore investigated whether such coarse-to-fine tuning dynamics occurs in L3 and L4 MUAe of mouse V1 and whether the size-tuned functional connections from L3 follow this dynamics as well.

We first investigated center size and SI dynamics for size-tuned MUAe in L4 and L3 relative to stimulus onset (Figure 5a). Both L4 and L3 MUAe showed an initial phase with rapidly decreasing center sizes and increasing SIs, followed by a phase with stable RF properties. A similar two-stage dynamics has previously been shown in cat LGN (Einevoll et al. 2011; Ruksenas et al. 2007). To better quantify the observed dynamics and test whether it held across mice, we investigated whether center sizes negatively correlated with SI in the initial phase, using linear mixed effects models with owing variable intercepts and slope across mice. This revealed a consistent inverse relationship between RF center size and SI for L4 MUAe (slope −0.014; F(1,371)=17.36, p<0.001) and L3 MUAe (slope −0.012; F(1,131)=10.04, p=0.002). These results demonstrate that a rapid sharpening of tuning-curve profiles in the first 150 ms occurs reliably across mice.

**Figure 5.**
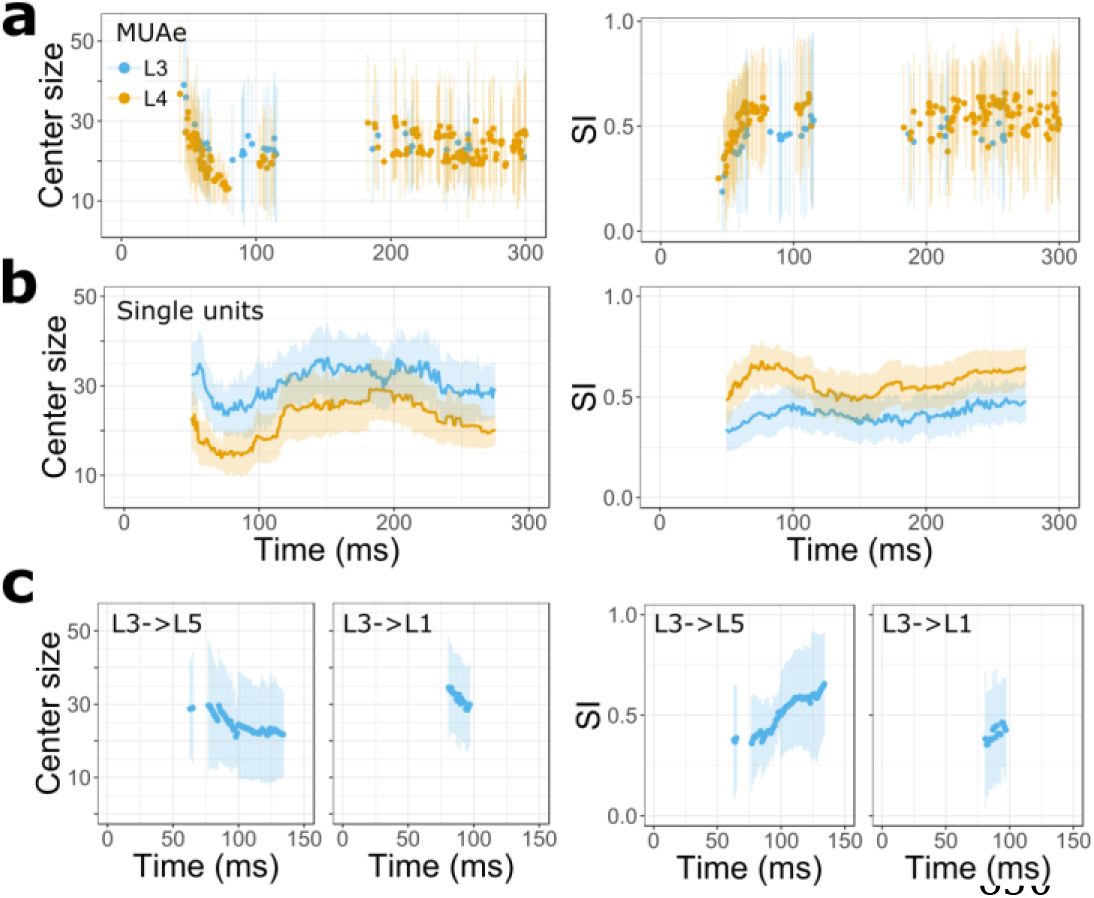
Dynamic sharpening of V1 RFs. a) Average center sizes (left) and SI (right) for size-tuned L4 and L3 MUAe. Lines reflect 95% confidence intervals across mice. b) Center sizes and SI across single-units in L3 (n=35) and L4 (n=29). Fits were performed for trial-averaged spike rates (bin size of 1ms) using a 50 ms sliding window. Error bars reflect 95% confidence intervals across neurons. c) Average SI and center sizes for functional connectivity from L3. Error bars reflect 95% confidence intervals.

We found similar coarse-to-fine tuning when we investigated RF dynamics of single-units (see Methods, Extracellular recordings). We identified single neurons in L3 and L4 that showed surround suppression (L3, n=35; L4, n=29) and used a moving window approach to determine their RF dynamics, obtaining good RoG model fits (Supplementary Figure 3). Inspecting RF center size and suppression strength dynamically in 50 ms moving windows, we found a sharpening of receptive fields between 50 and 100 ms after stimulus onset, with simultaneously decreasing center sizes and increasing suppression strength (Figure 5b). This RF sharpening observed in single-units provides a physiological basis for the sharpening seen in MUAe and functional connections, lending support to the notion that surround-suppressed MUAe and functional connections qualitatively reflect the underlying activity of single-units.

We finally inspected whether similar dynamics existed for the L3->L5 and L3->L1 connections, which are the most sustained of the surround-suppressed connections (Figure 4). We found that these connections showed similar tuning dynamics as observed in MUAe and single-units, with decreasing center size and increasing SI between 50 and 150 ms after stimulus onset (Figure 5c). Like for MUAe, center sizes negatively correlated with SI across animals for the L3->L5 (−0.03; F(1,397)=13.14, p<0.001) and L3->L1 connection (−0.02; F(1, 103)=47.06, p<0.001). This sharper tuning of functional connections proceeded in parallel with sharper tuning of MUAe activity in L3, suggesting that relays of size information to L5 and L1 qualitatively follow the coarse to fine dynamics observed in L3 MUAe.

It is remarkable that MUAe, single-units and functional connections showed similar coarse-to-fine dynamics of tuning parameters, particularly because functional connections were derived from low frequency LFP signals that reflect a complex mixture of cellular and postsynaptic currents (Buzsáki et al. 2012; Einevoll et al. 2013). A parsimonious interpretation of these converging findings is that there is a common local source for this sharpening in single-unit activity.

**Table 2.** RoG model descriptors per connection, median (min, max) across animals

|  | SI | Center size | R<sup>2</sup> | Frequency | log B<sub>12</sub> |
| --- | --- | --- | --- | --- | --- |
| L3->L1 | 0.3 (0.1-0.83) | 38.57 (15.9-43.98) | 0.81 (0.63-0.92) | $\beta, \gamma$ | 4 (2-9) |
| L3->L4 | 0.68 (0.45- 0.82) | 21.4 (11.17-26.73) | 0.61 (0.57-0.72) | $\gamma$ | 4 (2-6) |
| L3->L5 | 0.59 (0.11- 0.73) | 16.58 (10.15-49.4) | 0.85 (0.65-0.92) | $\beta, \alpha, \theta, \gamma$ | 8 (3-13) |
| L4->L2 | 0.65 (0.05- 0.89) | 23.09 (10.15-40.94) | 0.56 (0.53-0.86) | $\beta$ | 3 (2-5) |
| L6->L4 | 0.5 (0.35- 0.78) | 30.79 (18.95-36.2) | 0.65 (0.51-0.76) | $\gamma$ | 3 (2-4) |

## Discussion

We here provide a dynamic view on how spatial integration evolves across cortical layers of mouse V1 based on a Bayesian model comparison approach applied to laminar multi-unit activity and inter-laminar functional connectivity strengths. Our analyses reveal that information about stimulus size and context evolves across time and is dynamically communicated between cortical layers through a network of size-tuned functional connections (summarized in Figure 6). These connections, from L3, L4 and L6, parametrically vary with spatial context, driving activity in target layers L1, L2, L4 and L5 most strongly for intermediate stimulus sizes while showing reduced influence for larger ones. Amongst these functional connections, L3 occupies a central role, exhibiting surround suppression in its single- and multi-unit activity, as well as in its impact on other layers. These findings shed new light on how laminar activity is coordinated across a cortical column and on the different functional roles of cortical layers in spatial integration.

**Figure 6.**
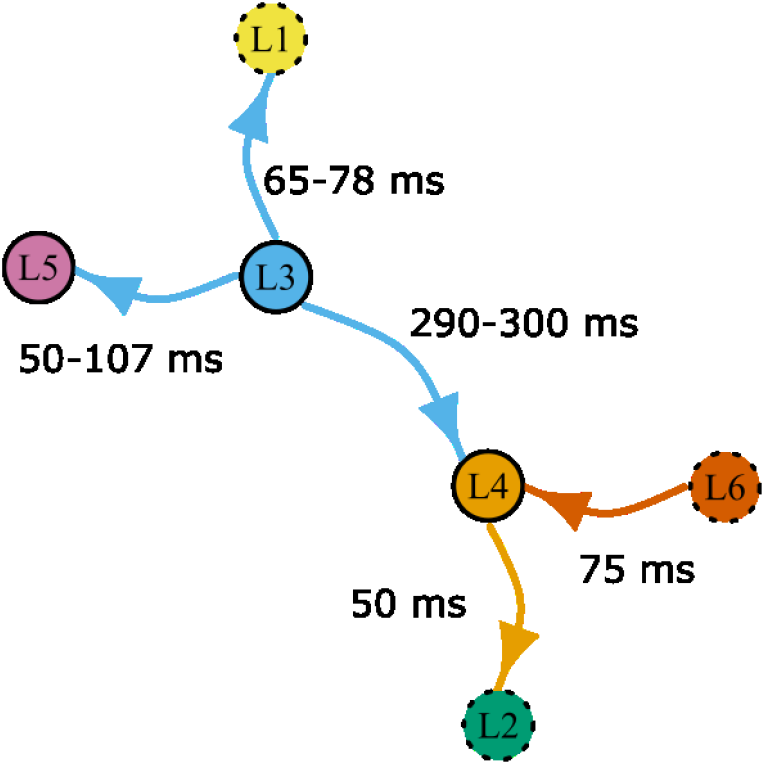
Summary of size-tuned functional connections obtained across time and frequencies. Each layer is considered a network node, and nodes are plotted closer together when a connection between them exhibits size tuning. Solid circles indicate layers at which surround-suppressed driving co-occurred with surround suppressed multi-unit activity in the source layer.

In line with previous anatomical and circuit-level results, our functional connectivity analyses reveal a major role for L3 in dynamically orchestrating spatial integration across cortical layers. In visual cortex of many mammalian species, L3 is well known for its prominent horizontal connectivity (Gilbert & Wiesel 1983; Rockland & Lund 1982), where neurons can extend their axons within the layer beyond their own RF. Being preferentially connected according to similarity in orientation preference (Ko et al. 2011; Bosking et al. 1997) makes these pyramidal cells optimally suited to mediate the well-known orientation dependency of surround modulations (Self et al. 2014; Nelson & Frost 1978). In addition, the preferential recruitment of SOM+ inhibitory interneurons by L2/3 pyramidal cells is a circuit motif in accordance with their prominent role in L2/3 surround suppression (Adesnik et al. 2012) or lateral inhibition (Pluta et al. 2017). Our finding of consistent and relatively strong surround suppression for considerable durations in L3 multi-unit and single-unit activity, are in line with this notion of a prominent role of L2/3 in shaping spatial integration.

In addition to exhibiting surround suppressed activity, our functional connectivity analyses also revealed that L3 coordinates activity in the column by modulating activity in L5, L1 and L4 according to spatial context. The predominant frequencies of L3 driving were in the beta and lower gamma band. Gamma band activity has been associated with feedforward streams at L3 (Bastos et al. 2015; Markov et al. 2013), speaking in favor of a feedforward interpretation. The L3->L5 connection also showed driving in lower bands, at longer latencies, that may play a role in feedback from downstream areas (von Stein & Sarnthein 2000). Generally, amongst the size-tuned connections from L3, the L3->L5 driving was most prominent, and could contribute to establishing surround-suppressed activity in L5, which occurred there at slightly longer latencies, and was notably briefer and weaker. Such a dual role of L3 in providing horizontal competition within L3, while driving a less suppressed signal in L5 is reminiscent of results in somatosensory cortex, where a layer specific excitation-inhibition ratio creates lateral suppression in L3 and feedforward facilitation in L5 (Adesnik & Scanziani 2010).

Remarkably, L5 surround-suppressed activity itself did not parametrically drive activity in other layers, even though L5 is known to play an important role in propagating activity across the column (Sakata & Harris 2009; Plomp et al. 2017) and sustaining L2/3 activity, including its horizontal spread (Wester & Contreras 2012). One would therefore hypothesize that L5, an output layer with cortico-cortical projections (Gilbert & Wiesel 1979; Harris & Shepherd 2015) and direct projections to subcortical regions (e.g. SC) and the contralateral hemisphere (Swadlow 1983; Kasper et al. 1994), has little role in providing size- and context-related information across the cortical column during spatial integration. Similarly, our results suggest that L4 plays a minor role in orchestrating activity in the column in a context-sensitive manner. L4 is a crucial relay of afferent activity to other layers, but with the exception of a brief functional connection to L2, L4 connections did not parametrically reflect stimulus context, even though L4 activity showed sustained periods of strong surround-suppression.

The parametric relay of size- and context information from L3 to L1 might help determine how feedback affects ongoing activity in the column. A preserved feature across mammals is that L1 contains relatively few cell bodies, local and lateral connections, but instead receives dense feedback projections (Thomson & Bannister 2003; Binzegger et al. 2004; D’Souza & Burkhalter 2017) and thalamic input (Harris & Shepherd 2015; Rockland 2017). L1 feedback is thought to be particularly important for far surround modulation (Angelucci & Bressloff 2006; Angelucci et al. 2017). L1 can modulate activity in supra- and infragranular layers through the apical dendrites of pyramidal cells (Egger et al. 2015; Jiang et al. 2013; Larkum 2013) and likely contributes to feedback-related enhancements of perceptual thresholds and discrimination performance (Takahashi et al. 2016; Zhang et al. 2014). Specifically, spatially specific top-down modulation via feedback to L1 might contribute to integration of information from outside the RF, such as occurring during contour integration and figure ground segregation (Self et al. 2013; Chen et al. 2017; Liang et al. 2017), and may also help support perceptual pop-out and segmentation of object boundaries (Angelucci et al. 2017; Coen-Cagli et al. 2012). We note, however, that in the absence of known L3->L1 excitatory projections, the observed functional connection could also be an indirect one, or result from a common top-down mechanism that manifests itself slightly earlier in L3 than L1. These hypotheses provide interesting directions for future investigations.

Our results are based on directed functional connectivity analysis within the Granger causality framework (Bressler & Seth 2011; Baccalá & Sameshima 2001; Granger 1969). The multivariate measure of Granger causality used here has a statistical interpretation of increased predictability between recorded signals that does justice to the directedness of neural interactions and accounts for their dynamics, but an interpretation in terms of neural circuits is not immediately warranted (Chen et al. 2017; Seth et al. 2015). While structural and functional connectivity are often closely related, the presence of a functional connection only indicates that activities systematically covary in time. Likewise, the presence of a structural connection only implies the potential for interaction (Battaglia et al. 2014), whose magnitude will depend on synaptic strength and other circuit-level forms of gating (Wang & Yang 2018). Although functional connectivity does not necessarily equal circuit connectivity, applying the Granger causality framework to the analysis of LFPs has previously helped understand the role of thalamocortical interactions in visual attention (Saalmann et al. 2012), interactions between cortical layers at rest and during stimulation (Bollimunta et al. 2008; Brovelli et al. 2004; Plomp, Quairiaux, Kiss, et al. 2014; Chen et al. 2017; Liang et al. 2017), as well as frequency-specific feedforward and feedback interactions between visual areas that are in excellent agreement with known anatomy (Bastos et al. 2015; van Kerkoerle et al. 2014; Michalareas et al. 2016). One possible limitation is that connectivity analyses based on LFP signals contain non-local signals through volume conduction: potentials measured at one place also reflect activity at more distant locations (Kajikawa & Schroeder 2011; Buzsáki et al. 2012). Although we addressed this by using bipolar signals and an orthogonalized derivation of connectivity strengths (Trongnetrpunya et al. 2015; Omidvarnia et al. 2014), bipolar LFPs still can reflect both local spiking activity and post-synaptic potential variations that may result from projections in the area. In our data, however, the fact that the most important surround-suppressed connections co-occurred with surround-suppressed multiunit activity gives credence to the interpretation that the firing of excitatory populations gives rise to directed functional interactions with a size-tuned profile. This was further corroborated by the similar dynamics of RF sharpening observed in functional connections as well as multi- and single-unit activity. Taken together our results provide a useful dynamic model of the interdependencies between activity measured at each layer of V1 and of how spatial context changes these relations.

In line with previous findings of receptive field dynamics in monkey and cat (Briggs & Usrey 2011; Malone et al. 2007; Wörgötter et al. 1998; Ruksenas et al. 2007), we show that in mouse V1 too the spatial receptive fields evolved from broad to sharp spatial tuning within the first 150 ms after stimulation. This dynamic sharpening was not only observed in L3 and L4 MUAe, but also in single-unit spike rates and functional connection strengths from L3 (Figure 5). The RF sharpening of activity in L3 seems to be independent of activity elsewhere in the column, because during this dynamic sharpening, size-specific connections did not target this layer. Hence, the RF sharpening might be related to known mechanisms within L3, such as surround suppression arising in supragranular layers through the inhibition of excitatory cells, via horizontal connections (Adesnik et al. 2012; Ozeki et al. 2009). What could be the role of such coarse-to-fine visual processing? The size of RFs is known to be contrast-dependent, such that the observed shrinkage over time suggests that the local processing in L3 and L4 might perform an enhancement of effective contrast, sharpening the representation of visual space. In line with this, the latencies match those of boundary detection processes, as observed in macaque V1 (Poort et al. 2016).

The fact that surround suppression is not restricted to V1 but present already at earlier processing stages raises the important question to which degree cortical surround suppression is inherited from dLGN or the retina. Neurons in mouse retina (Stone & Pinto 1993) and dLGN (Piscopo et al. 2013; Erisken et al. 2014), similarly to those in other mammals (Alitto & Usrey 2008; Jones et al. 2000; Solomon et al. 2002), have extraclassical suppressive surrounds, and geniculo-cortical afferents to V1 terminate not only in L4 but also target supragranular layers (Antonini et al. 1999; Cruz-Martín et al. 2014). Furthermore, orientation-specific surround modulations of L4 of mouse V1 do not seem to depend on activity in supragranular layers (Self et al. 2014). Together, this leaves open the possibility that - in addition to the prominent role of L3 found here - surround suppression in mouse V1 might also heavily depend on subcortical sources.

In the past, a number of studies have compared latencies of response onset with those of suppression onset to differentiate between inheritance and local mechanisms in surround suppression (Alitto & Usrey 2008). In our data, we observed delays of suppression relative to response onset of about 10 ms in L4 and L3. Similar delays, albeit of overall larger magnitude, between response onsets and suppression onsets have been previously reported in V1 of the anesthetized mouse (Self et al. 2014), and in macaque V1 for uniform patterns (Knierim & van Essen 1992; Smith et al. 2006). Other studies, however, have found approximately instantaneous onsets of suppression at response onset (Müller et al. 2003). In the meantime, it has become clear that the temporal evolution of surround modulation depends systematically on the strength of surround stimulation (Henry et al. 2013b). Since most of our main results rely on activity of local populations, which in rodents lack a large-scale organization according to preferred orientation (Ohki et al. 2005), our stimulus, at least on average, might indeed provide suboptimal drive to the extraclassical orientation-tuned surround mechanisms in V1. Together, the latency analysis and the prominent role of L3 in surround suppression observed here, therefore lend support to the interpretation that cortical mechanisms dynamically shape V1 spatial integration. We must note, however, that this evidence is not conclusive, because a delayed onset of suppression in V1 could, in principle, also be attributed to slowly developing signals inherited from dLGN or even the retina. Future studies performing a dynamic analysis of simultaneously recorded LGN and V1 activity and comparing the time course of surround suppression are clearly needed to unequivocally distinguish between these possibilities.

Probing MUAe activity time point by time point in the dynamic model comparison analysis, we did not find consistent evidence for surround-suppressed activity in L2 or L1. We attribute this lack of surround suppression to the stringent criteria of our dynamic analysis, which required that 6/7 animals show both positive evidence in favor of the RoG model and positive SI index at a single time point after stimulus onset. This is a conservative approach to identify which layers consistently exhibit size-tuning across animals that may exclude points that show more between-animal variability in time. Indeed, time-averaged MUAe in L1 and L2 did show a surround-suppressed activity profile (Figure 1c), consistent with previous mouse V1 studies (Nienborg et al. 2013; Self et al. 2014; Vaiceliunaite et al. 2013). Together, this discrepancy between the dynamic and time-averaged analysis shows that surround-suppressed activity in L2 and L1 is less consistently time-locked to stimulus onset than the suppression we dynamically observed in L3, L4 and L5. This in turn suggests that L1 and L2 serve less time-critical functions in spatial processing.

During our recordings, mice were awake and placed on an air-cushioned ball which allowed them to either sit or run, but to maximize the number of trials our analysis was performed irrespective of behavioral state. It is known that locomotion alters spatial integration in mouse area V1 (Ayaz et al. 2013) and dLGN (Erisken et al. 2014), increasing the RF center size and reducing surround suppression. In our experiments, however, effects of locomotion are unlikely to induce systematic biases, because bouts of locomotion occur spontaneously across the 11 randomly interleaved stimulus conditions. Having established the basic laminar profile of surround suppression within our dynamic connectivity analysis framework, it will be interesting to investigate the influence of behavioral state on laminar relay of size and context information in future studies.

Our results are limited to the ascending and descending relays of size information within a V1 column. In the future, it will be important to extend our recording approach to multishank probes and determine the relative contributions of vertical and horizontal interactions in V1 during spatial integration (Constantinople & Bruno 2013; Kätzel et al. 2011; Narayanan et al. 2015). Our analysis approach would also benefit from being applied to conditions with layer-specific or interneuron specific perturbations of neural circuits. Such causal manipulations would strongly constrain the functional connectivity models and provide further guidance in the interpretation of the model results.

## Acknowledgements

We thank Mattia F. Pagnotta for implementing row-normalized OPDC.

## Supplementary material

**Supplementary Figure 1.**
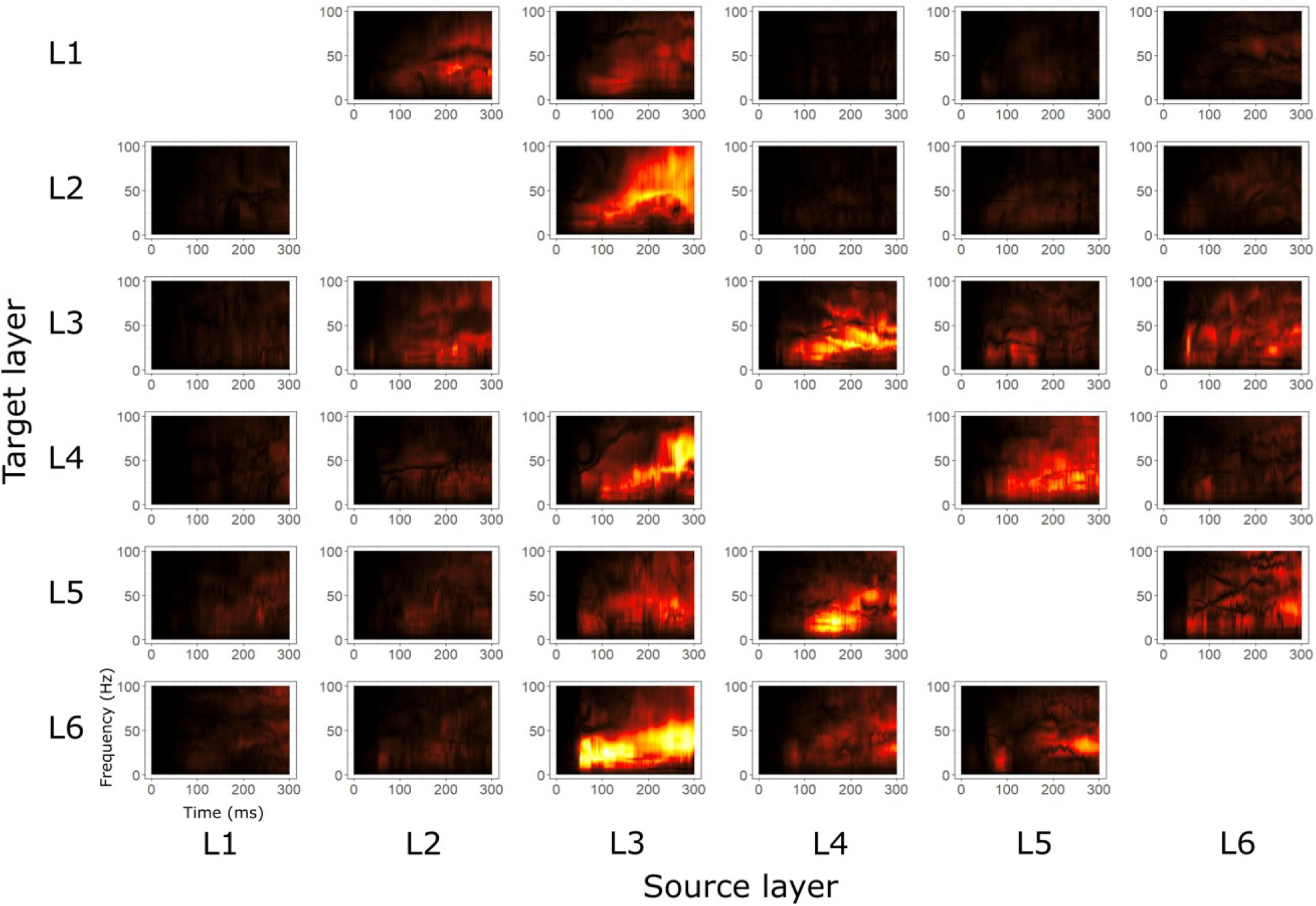
Time-frequency plots of directed functional connectivity strengths (median across animals) for each directed connection between all layers, in response to a 67° grating.

**Supplementary Figure 2.**
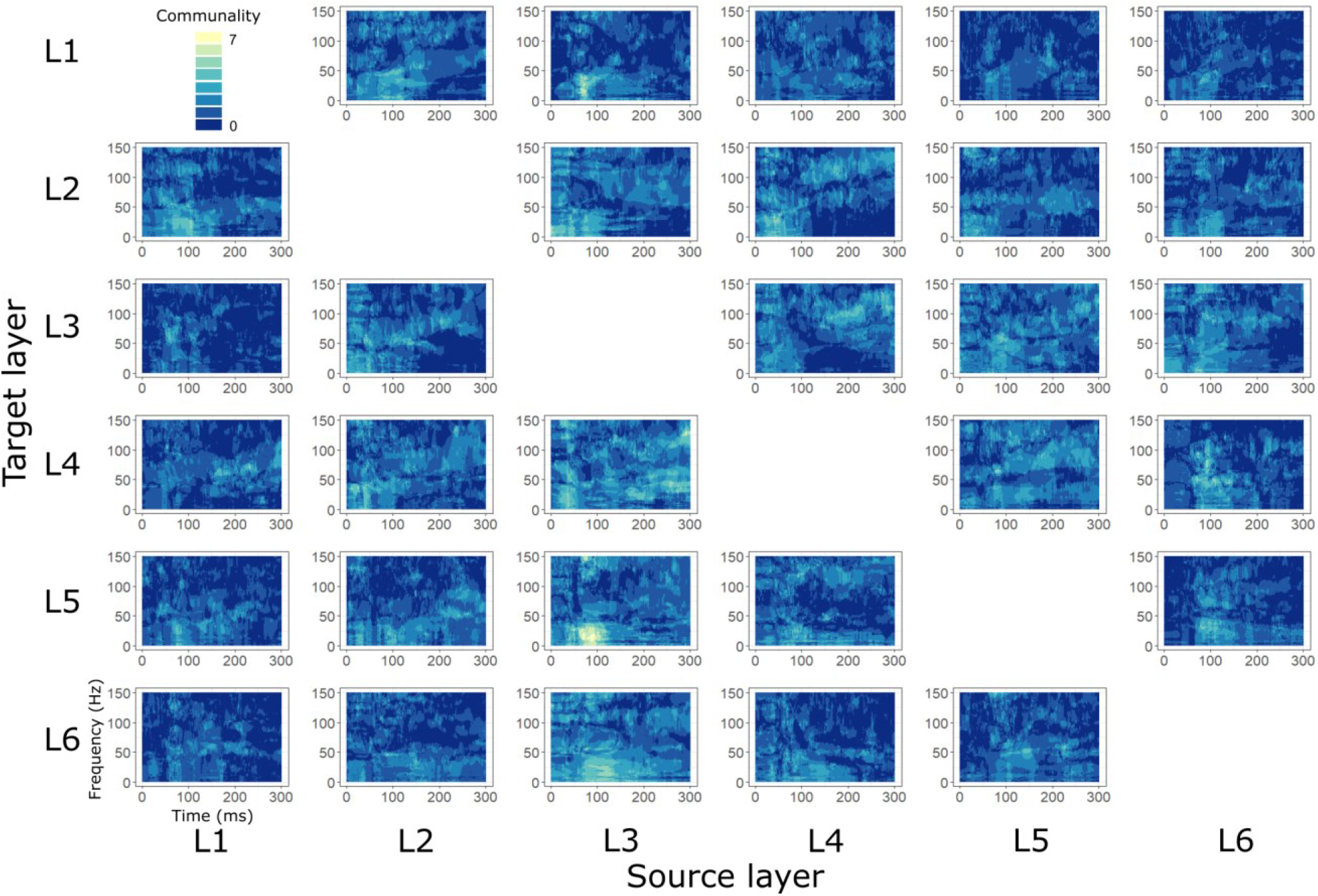
Time-frequency connectivity matrix showing the number of animals (0-7) passing both criteria (B_12_>3, SI>0), for the directed functional connection between all layers.

**Supplementary Figure 3.**
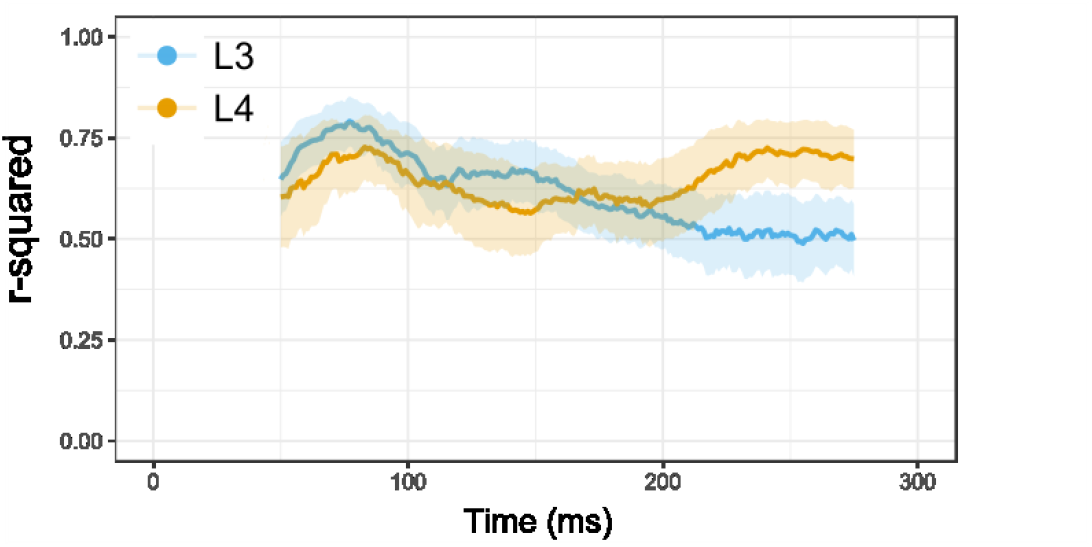
Dynamic single-unit model fits. Average R^2^ and 95% confidence intervals for RoG model fits of single-unit activity in the moving window analysis in L4 (n=29) and L3 (n=35).

